# Urolithin A induces cardioprotection and enhanced mitochondrial quality during natural aging and heart failure

**DOI:** 10.1101/2023.08.22.554375

**Authors:** S. Liu, J. Faitg, C Tissot, D. Konstantopoulos, R. Laws, G. Bourdier, P.A. Andreux, T. Davey, A. Singh, C. Rinsch, D.J. Marcinek, D. D’Amico

## Abstract

Cardiovascular diseases remain the primary cause of global mortality, necessitating effective strategies to alleviate their burden. Mitochondrial dysfunction is a driving force behind aging and chronic conditions, including heart disease. Here, we investigate the potential of Urolithin A (UA), a gut microbiome-derived postbiotic that enhances mitophagy, to ameliorate both age-related decline in cardiac function and cardiac failure. We highlight the significance of targeting mitochondria, by comparing gene expression changes in aging human hearts and cardiomyopathies. UA oral administration successfully counteracts mitochondrial and cardiac dysfunctions in preclinical models of aging and heart failure. UA improves both systolic and diastolic heart functions, distinguishing it from other mitochondrial interventions. In cardiomyocytes, UA recovers mitochondrial ultrastructural defects and decline in mitochondrial biomarkers occurring with aging and disease. These findings extend UA’s benefits to heart health, making UA a promising nutritional intervention to evaluate in the clinic to promote healthy cardiovascular function as we age.

## Introduction

Cardiovascular disease results in over $320 billion in public health spending per year^1^ and age is one of the greatest risk factors for cardiovascular disease^2^. Age-related changes in structure and function of the heart include thickening and stiffening of the left ventricular walls, increased fibrosis, and metabolic disruptions that lead to declines in cardiac function^3,4^, exercise intolerance and fatigue. This ultimately results in poor quality of life, loss of independence, and age-related comobidities^,5,6,7^. Although heart failure with preserved ejection fraction (HFpEF) is the predominant form of HF in older adults^8^, age-related comorbidities like hypertension are associated with reduced fractional shortening and contractility contributing to systolic dysfunction and increased sensitivity to additional cardiac insults^9,10^. Loss of skeletal muscle mass and a decline in VO_2_ are also highly correlated with HFpEF^11,12^,which further exacerbates the exercise intolerance and development of frailty.

Preclinical studies showed that mitochondrial dysfunction is a key driver of function decline in the aging heart. Changes in cardiac mitochondria include impaired ATP production, elevated oxidant production and a substrate shift from fatty acid toward glucose oxidation^13,14,15^. Given the high ATP demand of the heart, 90% of which goes to support Ca^2+^ pumping and cross-bridge cycling, the heart is especially sensitive to impaired mitochondrial energetics^16^. While impaired ATP production is a key factor linking aging mitochondria to cardiac dysfunction, mitochondrial dysfunction also disrupts cell signaling and protein post-translational modifications that can directly alter calcium dynamics and sarcomere function in the cardiac dysfunction^17,18,19^.

The accumulation of poorly functioning mitochondria with aging is associated with a deficit in mitochondrial quality control through mitophagy in multiple tissues, including the heart^20–22^. Therefore, targeting mitophagy using natural compounds is emerging as an exciting strategy for cardiac aging. Urolithin A (UA) is a naturally derived postbiotic produced by the gut microbiome from ellagitannin and ellagic acid compounds found in many fruits and nuts^23^. UA has been observed to activate mitophagy, mainly through both Pten induced Kinase 1 (PINK1)/Parkin dependent pathway and improve function in the skeletal muscles and joints in rodent models of aging and disease^24–26^. The effects of UA supplementation on cardiac function have translational potential. Three randomized clinical trials already showed that UA is safe, improved skeletal muscle endurance capacity in older adults^27^, and significantly increased muscle strength in overweight, middle-aged adults^28^, without exercise training.

Here, we investigated the role of mitochondrial and mitophagy dysfunctions in human cardiac aging and disease, and tested the hypothesis that UA supplementation reduces cardiac dysfunction in mice with age and in a heart failure model by improving mitochondrial function and mitophagy.

## Methods

### Human heart failure and GTEX RNA-seq data

Data from a human heart failure study were downloaded from the GEO database under the accession number GSE116250, using the fastq-dump command from the SRA toolkit. The data were downloaded in the form of single-end FastQ files, with 64 FastQ files corresponding to 64 samples. The data are summarized under three conditions: Non-failing (NF) with 14 samples, Dilated Cardiomyopathy (DCM) with 37 samples, and Ischemic Cardiomyopathy (ICM) with 13 samples. Heart data from GTEX were downloaded from the supplemental material of the study by Yang et al^29^, including information regarding upregulated and downregulated genes during aging, which correspond to positive and negative values of the "age" coefficient in the applied regression model, respectively. Additionally, normalized expression values, represented as Transcripts Per Million (TPM), for the respective RNA-seq samples (432 individuals across 54,592 genes) were obtained from the GTEx Consortium Portal (https://gtexportal.org/home/datasets). The dataset encapsulates six distinct age groups: 20-29 (22 samples), 30-39 (26 samples), 40-49 (66 samples), 50-59 (154 samples), 60-69 (150 samples), and 70-79 (14 samples). Data analysis was conducted using a linear regression model to detect aging-related genes. Genes were categorized as either "upregulated" or "downregulated" based on the sign of the "age" coefficient in the corresponding regression model (denoted as "Age coef"). In this study, this dataset is referred to as the "GTEx aging" dataset."

### Differential expression analysis of human heart failure data

Differential expression analysis was applied between “Ischemic Cardiomyopathy “(ICM) and “Non-failing” (NF) conditions (referred as “ICM_vs_NF”), and between “Dilated Cardiomyopathy” (DCM) and “NF” conditions, using “NF” as reference dataset (referred as “DCM_vs_NF”).

Before the DE analysis, a pre-filtering step was applied to remove low abundance mRNA measurements, by excluding genes with less than 10 total counts across samples. Additionally, an independent filtering procedure of DESeq2 was enabled, in order to filter out genes with very low counts that are unlikely to show significant alterations in gene expression. Gene symbols (MGI nomenclature), descriptions, and biotypes were matched to Ensembl GTF ids by using the R package biomaRt. v. 2.46.3. P-values were corrected for multiple-testing using the Benjamini and Hochberg (BH) method [Benjamini and Hochberg, 1995].

### Gene Set Enrichment Analysis (GSEA) – Human heart aging and heart failure datasets

The database used for GSEA enrichment was the Cellular Components collection of the Gene Ontology database (GO CCs). The enrichment analysis for the human heart aging (GTEx aging) and for each heart failure comparison (“ICM_vs_NF” and “DCM_vs_NF”) was performed separately, using the R package ClusterProfiler.

For the GTEx aging dataset, the age coefficient (“Age Coef”)^29^, was used as a gene ranking metric, while for each heart failure comparison, genes were ranked by the respective log2 fold change value. Consequently, the enrichment of GO CC gene sets among up- or-down regulated genes was statistically tested. The minimum and maximum gene set sizes were set to 10 and 500, respectively. The resulting p-values were adjusted using the BH method.

Terms with an adjusted p-value ≤ 0.05 were considered as statistically significant and were further characterized as activated or supressed based on the respective Normalized Enrichment Score (NES) sign (activated for NES > 0 and supressed for NES < 0). Common and unique repressed GSEA GO CCs between heart aging and heart failure datasets were visualized as a Venn diagram, using the R VennDiagram package. Common repressed terms between the GTEx aging and the “ICM_vs_NF” and “DCM_vs_NF” were further visualized as a dot plot using the enrichplot R package. The genes that commonly contributed to the enrichment of the “Mitochondrial inner membrane” GO CC term for the human heart failure comparisons were visualized as heatmaps of scaled normalized expression counts. Similarly, for the GTEx aging GSEA results, the core enrichment genes of “Mitochondrial inner membrane” GO CC term was visualized as a heatmap of scaled Transcript Per Million (TPM) counts across aging. Genes were hierarchically clustered with a Euclidean distance metric using complete linkage.

### Cardiac aging animal studies

For the mitochondrial morphology and activity animal study (Study 1), animal experiments were performed in accordance to European guidelines for care and were approved by the Animal Ethical Committee, the Higher Education and Research Ministry. Twenty-four male C57BL/6Rj mice aged 21 months old and eighteen male C57BL/6Rj mice aged 8 weeks old were purchased from Janvier Labs (France). Mice were collectively housed in cages in controlled room at 22°C and 12 h light/ dark cycle at Biomeostasis’ facilities. All animals were allowed ad libitium access to an irradiated normal chow diet (NC, irradiated pellet A04; SAFE, Villemoisson-sur-Orge, France) and to ultrapure and laboratory-grade acidified water (Aquavive®, INNOVIVE, France). For 8 weeks, 21-months-old and 8-week-old mice were treated by oral gavage, either with a vehicle solution (0.5% carboxymethylcellulose) (21 months vehicle, n=12 or 8 weeks vehicle n=9) or with UA at 50mg/ml (21 months UA, n=12 or 8 weeks UA, n=9). One day after the last dosing, overnight-fasted mice were anaesthetized by Isoflurane. The hearts were collected, weighed and the LV was separated from the rest of the heart. One part of LV was embedded in OCT following the protocol described in previous study. (Meng H. et al., 2014). The other parts of the LV were collected for electron microscopy.

For cardiac phenotyping study (Study 2), all animal experiments were approved by the University of Washington Institutional Animal Care and Use Committee. A total of 40 aged (24-26 month) C57BL/6J mice and 10 young (5 months) male mice were received from NIA. All mice were maintained at 21 °C and a 14/10 dark cycle at 30-70% humidity feeding standard chow (LabDiet PicoLab Rodent Diet 20) and water ad libitum before the experimental diet was administered. Animals were fed with control diet (Research Diets AIN93G) or diet supplemented with UA oral targeting 50mg/kg/day for 8 weeks.

### In situ Muscle Force

For study 2, maximal forces were performed as described^30^ [PMID 23692570] using an Aurora Scientific 305C servomotor (Aurora, Ontario, Canada) at baseline and after 7 weeks of treatment. Briefly, each mouse was anesthetized with isoflurane (4% for induction and ∼2% for maintenance) and laid on its side on a temperature-controlled platform maintained at 37°C. The right knee was clamped in place and the foot was secured to a footplate with the ankle positioned at 90°. The tibial nerve was stimulated with a Grass Instruments S88X stimulator (Astro-Med, Inc., West Warwick, Rhode Island, USA) at an optimal voltage (1.5 V) using percutaneous electrodes. Maximal tetanic torque was assessed by a force-frequency curve, where the muscle was stimulated every other minute at frequencies from 10 Hz to 200 Hz.

### Echocardiography

At baseline and after 8 weeks of treatment mice from study 2 were anesthetized with isoflurane (4% for induction and ∼2% for maintenance), and echocardiography was performed using The Vevo 3100 preclinical imaging system, Vevo Imaging Station, and MS400 probe from VisualSonics (Toronta,Canada). The mice were placed in a supine position and HR, respiration, core body temperature are monitored using Vevo Animal Monitoring system SM200 and Vevo Monitor App software from VisualSonics. Parasternal short axis view (PSAX) B-mode and M-Mode images were acquired. Echocardiography was analyzed using the Vevo LAB software. ECG and heart rate were monitored throughout the procedure with mouse heart rates being maintained in the range of 450-550 bpm at the low workload. To induce a high workload 3ug/g body weight dobutamine was injected to induce increased systolic function. High workload echocardiography performed once the heart rate increased ∼100bpm and remained stable. All protocols were approved by the University of Washington Institutional Animal care and use Committee. Systolic functions including LV mass, ejection fraction, fractional shortening are quantified at rest (low workload) and high workload (Stabilized post IP injection). Diastolic function was accessed by tissue velocity from tissue doppler apical four chamber view (detailed description described previously, Gavin, BioRxiv). In brief, an average of at least 3 individual cardiac cycles without a respiration are used to calculate peak early (E’) and peak late (A’) mitral valve anulus tissue velocity. The results were exported to Microsoft Excel and statistical analysis and graphing is performed using GraphPad Prism9 software.

### Euthanasia and tissue handling

Mice from study 2 were euthanized after 16 hours overnight fasting by cervical dislocation after endpoint measurement. Heart, muscle, brain, kidney and liver were immediately removed and weighed. A 2mm section was removed from ventricles for histology and the remaining tissue were snap frozen in liquid N2 to store for further analysis.

### Transmission electron microscopy

The tissues from mice in study 1 were slightly teased apart in 0.1 M Sorenson’s Phosphate buffer and cut into 1 mm^3^ pieces. Samples were immersion fixed in 2% glutaraldehyde with 0.1 M Sorenson’s buffer (pH 7.3) at 4 °C.

Tissues were postfixed in 1% osmium tetroxide (1 h), dehydrated in graded acetone (25%; 50%; 75%; 2 × 100%; 30 min each, RT) before being impregnated with increasing concentrations of epoxy resin (TAAB medium resin) in acetone (25%, 50%, 75%, 3 × 100%, all for 1 h each, RT). The samples were then embedded in 100% fresh resin and left to polymerise at 60 °C for a minimum of 24 h.

All resin blocks were trimmed using a razor blade to form a trapezoid block face. Sections were cut in a longitudinal or transverse orientation on an ultramicrotome^31^ using a diamond knife. Semithin sections (0.5 μm) were stained with toluidine blue and viewed on a light microscope (LM) to verify orientation of tissue. Ultrathin sections (70 nm) were then cut and picked up onto copper grids. Sections were stained with 1% uranyl acetate (30 min) and 3% lead citrate (7 min). All sections were examined using a HT7800 120kV TEM (Hitachi). Digital micrographs were captured using an EMSIS Xarosa CMOS Camera with Radius software.

### Morphological and statistical analyses from TEM images

Mitochondrial shape descriptors and size measurements were obtained using Image J (version 1.52i, National Institutes of Health, Bethesda, MD, USA) by manually tracing mitochondria from TEM images. Surface area (mitochondrial size) is reported in squared micrometres; perimeter in micrometers; Values were imported into Microsoft Excel and Prism 8 software (GraphPad Software, San Diego, CA, USA) for data analysis. Statistical significance was evaluated based on 99% confidence interval (CI) of the median.

### Succinate Dehydrogenase activity

Hearts in OCT were cut into sections of 8 µM using a cryostat. Sections were then incubated in a solution containing nitroblue tetrazolium (1.5 mM), sodium succinate (130 mM), phenazine methosulphate (0.2 mM), and sodium azide (0.1 mM) for 6 min at room temperature. The reaction was stopped by two washes in PBS. Slides were mounted using Fluoromont mounting medium (Thermofisher, 00-4958-02). All samples were processed successively using the same incubation solution ensuring that they underwent the exact same experimental conditions. Images of the tissues were taken with a bright field microscope. The intensity of the staining was analyzed with Image J.

### RNA extraction from mouse heart OCT samples

Excess OCT was trimmed away from the frozen tissues block with a sterile scalpel blade. The tissues were cut into sections of 20 µM using a cryostat previously cleaned with a RNAse away solution. 10 sections per sample were collected in microtubes containing 1mL of Trizol. The RNA was isolated using the TRIZOL/Chloroform extraction method.

### Mouse heart aging RNA-seq data

RNA was quality controlled with Agilent Fragment Analyzer System. Library preparation was performed by Strand-specific cDNA library, purification of poly-A containing mRNA molecules, mRNA fragmentation, random primed cDNA synthesis (strand-specific), adapter ligation and adapter specific PCR amplification. RNA-seq run was performed with the NovaSeq6000 using S4 flowcells with 2x150bp (>30 million read pairs (+/- 3%) per sample).

Data were grouped into four categories based on experimental conditions: Young with 4 samples, Young treated with UA (Young_UA) with 5 samples, Old with 6 samples, and Old treated with UA (Old_UA) with 3 samples. Quality Control (QC) was performed as described for the human heart failure study above. High quality RNA-seq read pairs were mapped to the Ensembl mouse GRCm38 reference genome, using a transcriptome annotation GTF file (version GRCm38.94). An in-house RNA-sequencing alignment pipeline that utilizes the STAR aligner version 2.6.1c was applied. For the alignment process, a bespoke index was built using the annotation file, and an expected read length of 151 bps. Read counting was performed on the gene-level, using htseq-count. To inspect the alignment results, MultiQC version 1.8.dev0 was applied. Read counts were normalised using the variance stabilizing transformation (VST) method, implemented in DESeq2. Principal Component Analysis (PCA) was performed using the normalised read counts of the 1,000 most variable genes (mvgs) using base R (prcomp() PCA function), and visualized using the R package ggplot2. Differential expression analysis was applied between “Old” and “Young” conditions, using “Young” as reference dataset (referred as “Old vs young”), and between “Old_UA” and “Old” conditions, using “Old” as reference dataset (referred as “Old UA vs Old”). Before the DE analysis, a pre-filtering step was applied to remove low abundance mRNA measurements, by excluding genes with less than 10 total counts across samples. Additionally, an independent filtering procedure of DESeq2 was enabled, to filter out genes with very low counts that are unlikely to show significant alterations in gene expression. Gene symbols (MGI nomenclature), descriptions, and biotypes were matched to Ensembl GTF ids by using the R package biomaRt. v. 2.46. P-values were corrected for multiple-testing using the Benjamini and Hochberg (BH) method [Benjamini and Hochberg, 1995].

Gene Ontology Cellular Components (GO CCs) database was used for GSEA enrichment of the mouse heart aging and rat heart failure datasets. The enrichment analysis was performed for each DEA comparison separately, using the R package ClusterProfiler. For each comparison, the gene’s log2 fold change was multiplied by the respective p-value (log2(Fold Change) * -log10(p-value)), and genes were ranked by the resulting value. Consequently, the enrichment of GO CCs gene sets among up- or-down regulated genes was statistically tested. The minimum and maximum gene set sizes were set to 10 and 500, respectively. The resulting p-values were adjusted using the BH method. Terms with an adjusted p-value ≤ 0.05 were considered as statistically significant and were further characterized as activated or suppressed based on the respective Normalized Enrichment Score (NES) sign (activated for NES > 0 and suppressed for NES < 0). For the mouse heart aging dataset, activated GSEA GO CCs of the “Old UA vs old” comparison were further filtered, keeping only the terms that were also significantly suppressed in the “Old vs young” comparison.

### Mitochondrial respiration assay

Isolated cardiac mitochondria were tested for respiration using Oxygraph 2k dual respirometer (Oroboros Instruments, Innsbruck, Austria). For the substrate-uncoupler-inhibitor titration (SUIT) protocol, 10 μM cytochrome c was added to each chamber to allow measurement of respiration in isolated mitochondria without limitation by membrane damage occurring during isolation. Approximately 100 μg heart mitochondrial homogenate was added to each 2 mL chamber. State three fatty acid oxidation was measured by addition PC (palmitoyl carnitine as substrate (Fig S3A). State three CI&CII respiration was stimulated by adding CI (Pyruvate, Malate, Glutamate) (Figure S3B) prior to stimulation with ADP followed by succinate for State 3 respiration (Figure S3C).

### Heart failure animal study

The experimental procedures were carried out in accordance with European guidelines for the care and use of laboratory animals (Directive 2010/63/UE). The protocol that was used to induce myocardial infarction (MI) in rats was approved by an Animal Ethical Committee (French National Committee N°71) and by the Higher Education and Research Ministry (#30207-2021021117467048 v4) on April 11th, 2021. Fifty Wistar male rats were purchased from Janvier Labs (France). Animals were collectively housed in cages at SYNCOROSOME’s facilities and had free access to food (RM1, SDS Dietex) and drinking water *ad libitium.* Surgery was initiated in 7-week-old rats that were separated into three groups. MI was induced in animals from Surgery Vehicle (n=20) and Surgery UA (n=20), by chronic left anterior descending coronary artery (LAD) ligation performed at day 0. Rats were anesthetized by an intraperitoneal injection (IP) of Medetomidine (0.5 mg/kg) and Ketamine (50mg/kg). The animals were then intubated and ventilated at 10 ml/kg tidal volume and 70-80 cycles/minutes. The body temperature was maintained around 37°C using a thermoregulated heating pad connected to a rectal probe. Rats were placed in supine position, chest-shaved and prepared for standard surgical aseptic conditions. A left lateral thoracotomy was carefully performed to expose the heart. Suture was placed around the LAD (4/0 Silk, Ethicon), around 2 mm below the left atrium and close to the interventricular junction, to obtain infarct size (IS) near 40% of the total left ventricular (LV) area. The chest was closed, air was expelled from the rib cage to avoid pneumothorax and quick reanimation was performed using atipamezole hydrochloride (IM, 0,5 mg/ml, Zoetis).

The Sham Vehicle animal (n=10) were subjected to the same protocol. However, after the left lateral thoracotomy exposing the heart, the rib cage was closed without passing the suture thread around LAD. A peri-operative care of the animals was performed during the surgery.

For 8 weeks, starting 24 hours after the Surgery UA animals were daily treated with Urolithin A at 50 mg/ml by oral gavage. The rats from Sham vehicle and Surgery Vehicle group received the same dose of a vehicle solution (0.5% carboxymethylcellulose) at the same time. The volume of administration was adjusted every week based on the mean body weight.

### Echocardiography of heart failure study

To assess the cardiac remodeling and function following the MI, three echocardiography were performed 4 days, 1 month and 2 months after starting treatment. The first examinations were used to exclude rats with small infarcts and limited reduction of the ejection fraction.

Rats were anesthetized with 4% isoflurane in 50% oxygen-50% air mixture and maintained with 2-2.5% isoflurane during the procedure. The animals were placed in a supine position on a heating pad, thorax-shaved and imaged using a digital ultrasound system (Vivid 7,GE Medical Systems) equipped with a 10-Mhz phased-array and 13Mhz linear-array transducer. Standard B-mode (Brightness-mode) and M-mode (Motion-mode) images of the heart were obtained in the two-dimensional (2D) parasternal long axis view (PLAX). Fractional shortening and ejection fraction were calculated using the LV dimensions parameters previously described and the following equations:

FS = fractional shortening = [(LVIDd-LVIDs)/LVIDd] x 100.

EF= ejection fraction = [(end diastolic volume – end systolic volume)/ end diastolic volume]*100. The isovolumic relaxation time, ratio of peak velocity of early (E) and late (A) waves of the mitral flow during diastole and cardiac output were calculated by Doppler-echocardiography only for the last examinations. All parameters were measured and averaged from three consecutive cardiac cycles under stable conditions by a single blinded trained operator.

### Rat heart collection

One day after the last dosing, overnight-fasted rats were anesthetized with 2.5% isoflurane in 50% O_2_ 50% air mixture followed by an injection of buprenorphine (0.03mg/kg). Hearts were stopped in diastole (KCL 2M, 1ml/kg) by jugular injections. The atria were removed, the hearts were washed in cold PBS and quickly sponged to remove the liquid from the ventricles. LV was separated from the remaining part of the heart, weighed and incubated in NBF 10% for 48h. Hearts were transferred in ethanol 70%, cut in three transverse sections and embedded in paraffin in three different blocks, deriving from apical, middle and basal LV regions (Group 1, N=10, group2 and N=20).

### Infarct area and fibrosis quantification

Briefly, tissues block from the apical, middle and basal LV regions were cut into sections of 5 µM using a microtomes. Sections were stained with Sirus-RED (SR). To evaluate the extent of interstitial fibrosis, the SR area was quantified with the digital image analysis Visiopharm software. The infarct area was measured using Image J and averaged per animal.

### Rat heart RNA extraction

The protocol was performed according to the manufacture instructions using the kit RNeasy FFPE (Qiagen, 73504). Tissues block were cut into sections of 20 µM and collected in microtubes. Samples were incubated for 3 minutes at 56°C in a deparaffinization solution (Qiagen, 19093). A step of digestion with Proteinase K was performed followed by a treatment with DNAses. RNA was purified using the RNeasy MinElute column and eluted in 30uL of RNAse free water.

### Rat heart RNA-seq

Data were grouped into three categories based on experimental conditions: Vehicle (Sham) with 4 samples, MI with 4 samples, and MI+UA with 5 samples. QC was performed as in the mouse RNA-seq study. High quality RNA-seq read pairs were mapped to the Ensembl rat mRatBN7.2 reference genome using a transcriptome annotation GTF file (version mRatBN7.2.108). PCA was performed as in the muse RNA-seq study and highlighted a clear outlier “MI_UA” condition) which was excluded from downstream analysis (visualization not shown). Differential expression analysis was applied between “MI” and “Sham” conditions, using “Sham” as reference dataset (referred as “MI vs sham”), and between “MI+UA” and “MI” conditions, using “MI” as reference dataset (referred as “MI UA vs MI”). DEA and GSEA were performed as described in the human RNA-seq section (see above). RNA-seq was performed as in the mouse RNA-seq study, with the exception of applying rRNA depletion instead of poly-A containing mRNA purification.

### Heart Immunohistochemistry

Tissue blocks from the middle LV regions were cut into sections of 4 µM using a microtome. Sections were deparaffinized in xylene, rehydrated in descending alcohol baths and brought into distilled water. Permeabilization was performed by 15 minutes incubation in 0.1% Triton X-100 at 37°C. Slides were washed with PBS, containing 0.1% Triton X-100 and blocked 1 hour in 5% goat serum (Jackson Immuno Research, 005-000-121) 3% BSA (Pan Biotech, P06-1391100). Rabbit anti-phospho-ubiquitin, (Millipore, ABS1513-and mouse anti-VDAC (Abcam, ab14734) were diluted to 1/50 and 1/500 respectively before incubation at 4°C overnight. Sections were washed in TBST and incubated for 2hours with the secondary antibodies: Alexa 555 goat anti-rabbit (Invitrogen, A21428) and Alexa 488 goat anti-mouse (Invitrogen, A11001). Slides were mounted with Vectashield Antifade Mounting Medium (Adipogen, H-1500). Images were acquired using a confocal microscope (Zeiss, LSM-880).

Images were analyzed using Image J. Background was removed by applying an intensity threshold. After converting the images in mask, particles were quantified after filtering to remove non-specific signal determined from negative controls.

### Statistical analysis

Statistically significant differences between 2 independent groups were determined with Student’s unpaired t-test. Unpaired t-test with Welch’s correction were used to determine significance for pairs (p < 0.05). Statistically significant differences between three independent groups were determined by one-way ANOVA test followed by Tukey’s multiple comparison test. For group analysis involving two variables, a two-way ANOVA followed by Tukey’s multiple comparison test was performed. P values less than 0.05 were considered significant. Statistical analysis of RNA-seq experiments is described in the corresponding sections.

## Results

### Mitochondrial Dysfunction is a Common Feature of Heart Aging and Disease in Humans

We investigated common signatures that characterize both cardiac aging and heart diseases. The study involved analyzing transcriptomics dataset from the human GTEx study, which included healthy subjects spanning a wide age range^32^. We compared this with another dataset analyzing transcriptional changes in patients with dilated cardiomyopathy (DCM) and ischemic cardiomyopathy (ICM), compared to non-failing healthy controls (NF)^33^. By applying gene set enrichment analysis (GSEA) and dataset integration, five gene sets were found to be commonly downregulated in both cardiac aging and in disease states (Fig. 1A and Suppl. Table 1). These gene sets encompassed mitochondrial categories, as well as “microtubule, “nucleoid” and “organellar ribosome” pathways (Fig. 1B). Interestingly, even the latter two gene sets included numerous mitochondrial genes, particularly those involved in mitochondrial transcriptional regulators (Suppl. Fig. 1A, B) and mitochondrial translation (Suppl. Fig. 1C, D). The pathways “mitochondrial matrix” (Suppl. Table 1) and “mitochondrial inner membrane” (Fig. 1C, D, Suppl. Table 1) exhibited the largest number of enriched core genes across datasets. The most significant downregulation was observed in genes involved in mitochondrial respiration and translation (Fig. 1E). Notably, we also observed significant downregulation of several mitophagy genes, including *PINK1* and *OPTN1*, as well as mitochondrial dynamics and cristae regulators, such as *OPA1* (Fig. 1F). This suggests that a decline in both mitochondrial function and mitophagy might be a common feature of human heart aging and cardiac diseases and indicate the potential to improve heart health using interventions promoting mitochondrial quality control pathways.

**Figure 1.**
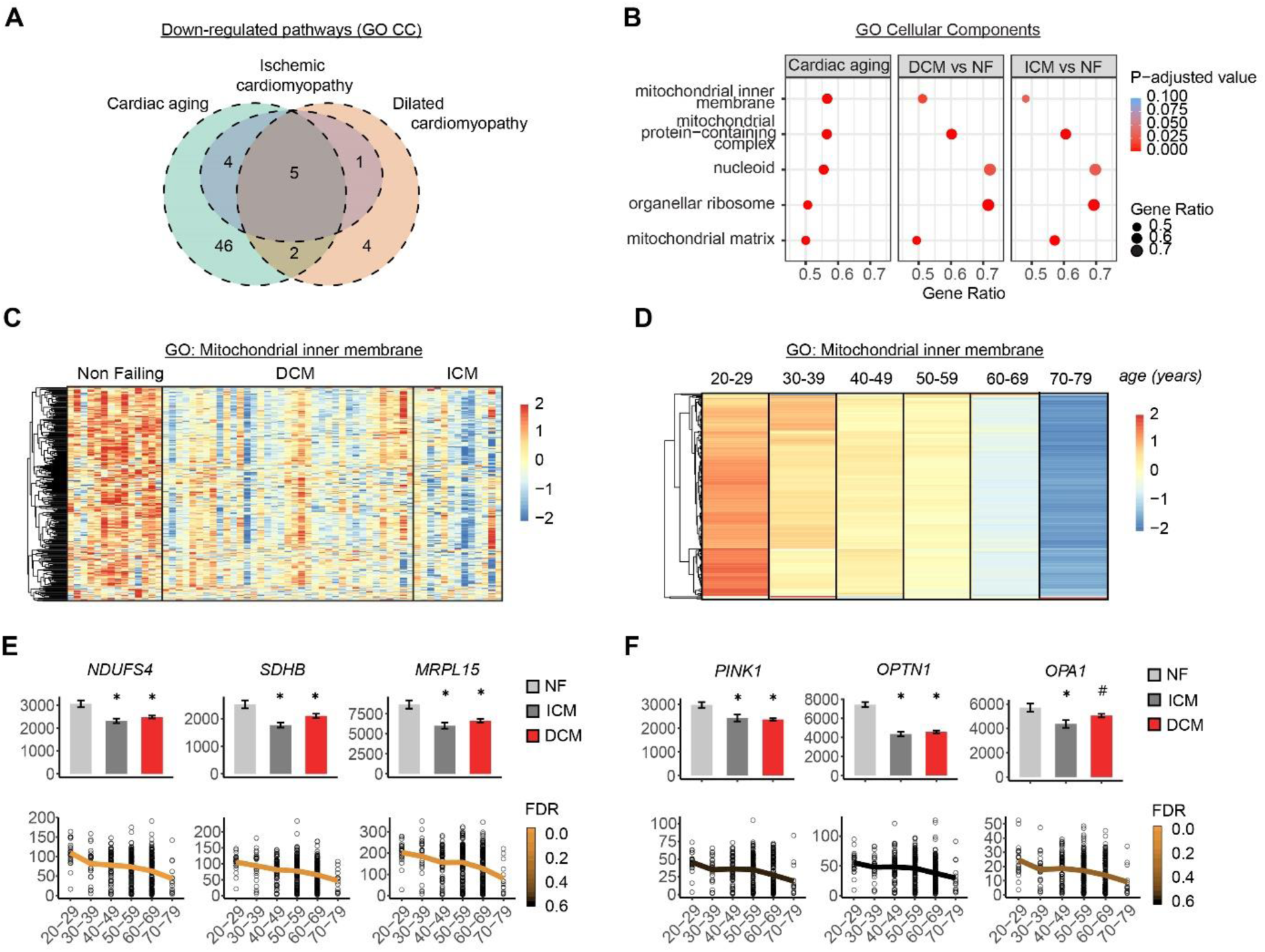
Mitochondrial dysfunction is a common hallmark of human cardiac aging and disease. A) Venn diagram depicting overlap of significantly down-regulated pathways (Gene Ontology Cellular Components, GO CCs) among three human datasets: Cardiac Aging, Dilated Cardiomyopathy (DCM) and Ischemic Cardiomyopathy (ICM). GO CCs were identified using Gene Set Enrichment Analysis (GSEA) and categorized as "down-regulated" based on their normalized enrichment scores. B) Dot plot of five commonly suppressed GO CCs shared among all three datasets in (A). GO CCs are sorted by "Cardiac Aging" Gene ratio. The size of each dot corresponds to the Gene Ratio, while the color indicates the statistical significance of the enrichment (Benjamini-Hochberg-adj. p-value). C) Heatmaps of scaled normalized expression counts (z-scores) of genes that contributed to the enrichment of the "Mitochondrial Inner Membrane" GO CC term in human DCM and ICM compared to Non-Failing. Rows represent genes hierarchically clustered using a Euclidean distance metric and complete linkage. Columns represent individual samples from each condition. D) Heatmaps of the scaled Transcripts Per Million (TPM, as z-scores) of core enrichment genes contributing to the "Mitochondrial Inner Membrane" GO CC term in the human cardiac aging study (GTEx Aging). Rows represent genes hierarchically clustered using a Euclidean distance metric and complete linkage. Columns represent different age groups, sorted in ascending order. For each age group, mean TPM values were calculated for all genes before scaling. E) Expression levels of top three most significantly downregulated genes in the “Mitochondrial inner membrane” category comparing cardiac diseases (ICM and DCM) to controls (NF) (bar plot, top) or comparing age groups (scatter plot, bottom). F) Expression levels of downregulated genes related to mitophagy (*PINK1*, *OPTN1*) and mitochondrial quality (*OPA1*). Comparisons as in (E). Bar plots illustrating normalized expression values. * for p < 0.05, # for p >= 0.05. Lower panel: Scatter plots showing Transcripts Per Million (TPM) expression. Mean TPM values were calculated for each age group and connected with a line, displayed in ascending order of age groups. The color scale of the line represents the level of significance (False Discovery Rate, FDR).

### Urolithin A Improved Heart Muscle Mitochondrial Morphology, Ultrastructure, and Function

We confirmed the impairment of mitochondrial morphology and ultrastructure during heart aging and the potential of mitophagy activators to revert this defect in a rodent model of natural aging. For this, we quantified mitochondrial ultrastructure (cristae) by transmission electron microscopy (TEM) in wild type mice aged 24 months (old) or 8 weeks (young) administered with either vehicle or the mitophagy inducer urolithin A (UA), for 2 months. Density and number of mitochondrial cristae was markedly decreased with age, as evidenced by the loss of strict parallel position (Fig. 2A and 2B). UA intervention rescued the age-dependent decline in cristae volume density to levels compared to those observed in young animals (Fig. 2A-B). Interestingly, mitochondrial cristae were more circular with age, while UA administration to old animals reverted such increase (Fig. 2C). Regarding mitochondrial morphology, mitochondria were smaller in old compared to young animals, an effect rescued by UA administration (Fig 2D). No differences in mitochondrial volume density and mitochondrial number were observed with UA in either young or old mice (Suppl. Fig. 2A and 2B). These results indicate that age-associated alteration in heart mitochondrial morphology and ultrastructure are reverted by 2 months of UA supplementation, while neither aging nor UA affected mitochondrial content.

**Figure 2:**
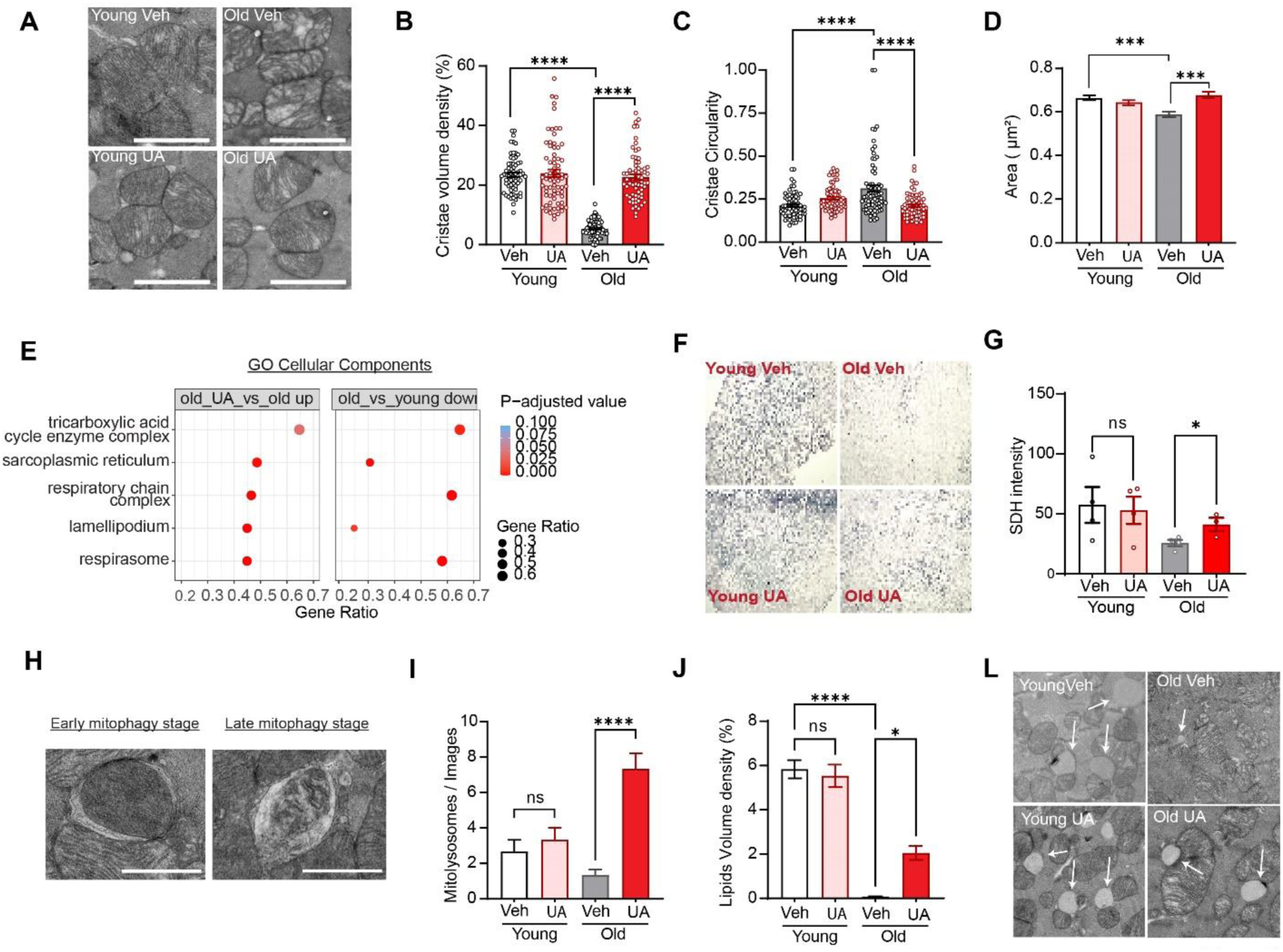
UA improves heart mitochondrial ultrastructure, morphology and function in old mice. A-C) Representative heart transmission electron microscopy (TEM) of transversal images of young and old mice administered with either vehicle or UA for 2 months (A) and corresponding quantification of mitochondrial cristae density (B) and circularity (C). N= 75 mitochondria /group. D) Quantification of mitochondrial area in transversal orientation in young and old mice from vehicle and UA groups (N=3 animal/group). E) Gene set enrichment analysis from house heart aging RNA-seq data describing Gene Ontology (GO) Cellular Component gene sets that are significantly repressed with cardiac aging (old vs. young down) and significantly rescued by UA administration to old mice (old UA vs old up). The size of each dot corresponds to the Gene Ratio, while the color indicates the statistical significance of the enrichment (Benjamini-Hochberg-adj. p-value). F-G) Representative succinate dehydrogenase (SDH) stain (F) and corresponding quantification G) of experimental groups as in Fig. 2A (N=4-3 animal/group). Scale bars: 5mM. G-H) Representative images of mitolysosome events (G) and their quantification in young and old mice treated as indicated (H) (N=3 animal/group). Scale bars: 500nm. J-L) Quantification of lipid droplet volume density expressed as percentage over total area in the indicated animals (J) and representative images with arrows pointing to lipid droplets (L). p value. *p<0.05; *** p<0.005; ****p < 0.001 after one-way ANOVA. Error bars represent mean ± SEM.

Heart RNA-seq in these animals further showed that the age-related decline in pathways associated with mitochondrial function is rescued by UA supplementation (Fig. 2E and Suppl. Table 2). To confirm the impact of UA on mitochondrial function, we measured the activity of the enzyme succinate dehydrogenase (SDH) in situ on heart muscle cross-sections (Fig. 2F, 2G and Suppl. Fig. 2C). A trend for a decrease in SDH activity was quantified in old compared to young mice. In contrast, SDH activity in old UA-treated animals was significantly higher than in old controls and to similar in levels to young animals (Fig. 2F, 2G and Suppl. Fig. 2C). Consistent with the human data above (Fig. 1F), aged mice showed a significant reduction in mitophagy events in the heart. UA supplementation significantly rescued the age-associated decline in mitophagy events (Fig. 2H-I). Finally, we quantified the volume density of heart lipid droplets and identified a dramatic reduction in old compared to young animals, and a partial but significant rescue in old mice treated with UA (Fig. 2J and 2L). These results indicate that 2 months of UA administration activate mitochondria recycling and renewing to achieve a mitochondrial pool with improved function.

### UA Improved Cardiac and Skeletal Muscle Function in Old Mice

After obtaining these positive biomarker data, we proceeded with performing functional cardiac measurements in 24-months old mice fed with urolithin A or standard chow diet for 2 months, using 8 weeks old young mice as controls. (Fig. 3A). To assess diastolic function, the ratio of blood flow across the mitral valve in early diastole (Ea/E’) to the flow in late diastole (Aa/A’) was determined. This was done using echocardiographic imaging, as illustrated in Figure 3B. A comparison between old and young mice revealed a reduced ratio of early to late diastolic mitral annulus velocities (Ea/Aa) in the aged mice, indicating a decline in diastolic function. Notably, the treatment of UA significantly preserved the age-related decline in diastolic function when compared to the old control group at the end of the 8-week treatment period (Figure 3C). When looking at the delta of E’/A’, comparing end of the study to baseline, the comparison between old UA and old control mice approached significance (P=0.0578) (Figure 3D). To assess systolic function, the percentage of fractional shortening (FS) was measured using echocardiography’s M-mode short axis. Anesthetized mice were subjected to lower-than-physiological heart rates, which may not accurately represent systolic function under compromised low workload conditions. Therefore, fractional shortening was also measured following the injection of the inotropic agent dobutamine to induce a higher heart rate within the physiological range (Fig. 3E). There was no significant difference in fractional shortening between groups at the low workload (LWL) (Fig. 3F). However, under higher workload conditions (HWL), UA treatment rescued the decline in fractional shortening associated with aging (Fig. 3F). Furthermore, a trend towards the preservation of ejection fraction (EF) was observed after 8 weeks of UA feeding (Fig. 3G). These changes in EF were statistically significant when analyzed as percentage comparing end of the study to baseline between the old UA-treated and old groups (Fig. 3H). Increased left ventricular mass (LVM) was observed with aging and suggests cardiac hypertrophy, either due to increased functional demand or pathological conditions specific to the heart or in combination with other organs. UA treatment only mildly affected LV mass hypotrophy phenotype in aged mice (Figure 3I). Functional changes in skeletal muscle were assessed after 8 weeks of UA feeding. The maximal torque of the plantar flexors was tested comparing end of the experiment to baseline for both control and UA-treated old mice. The UA group maintained maximal force compared to baseline, an effect which was significantly different when compared to the decline over time observed in the old control group (Fig. 3J). We also tested UA treatment on heart mitochondria respiration by oxygraphy. No difference was detected between the old controls and the old UA-supplemented group (Suppl. Fig. 3A-C).

**Figure 3.**
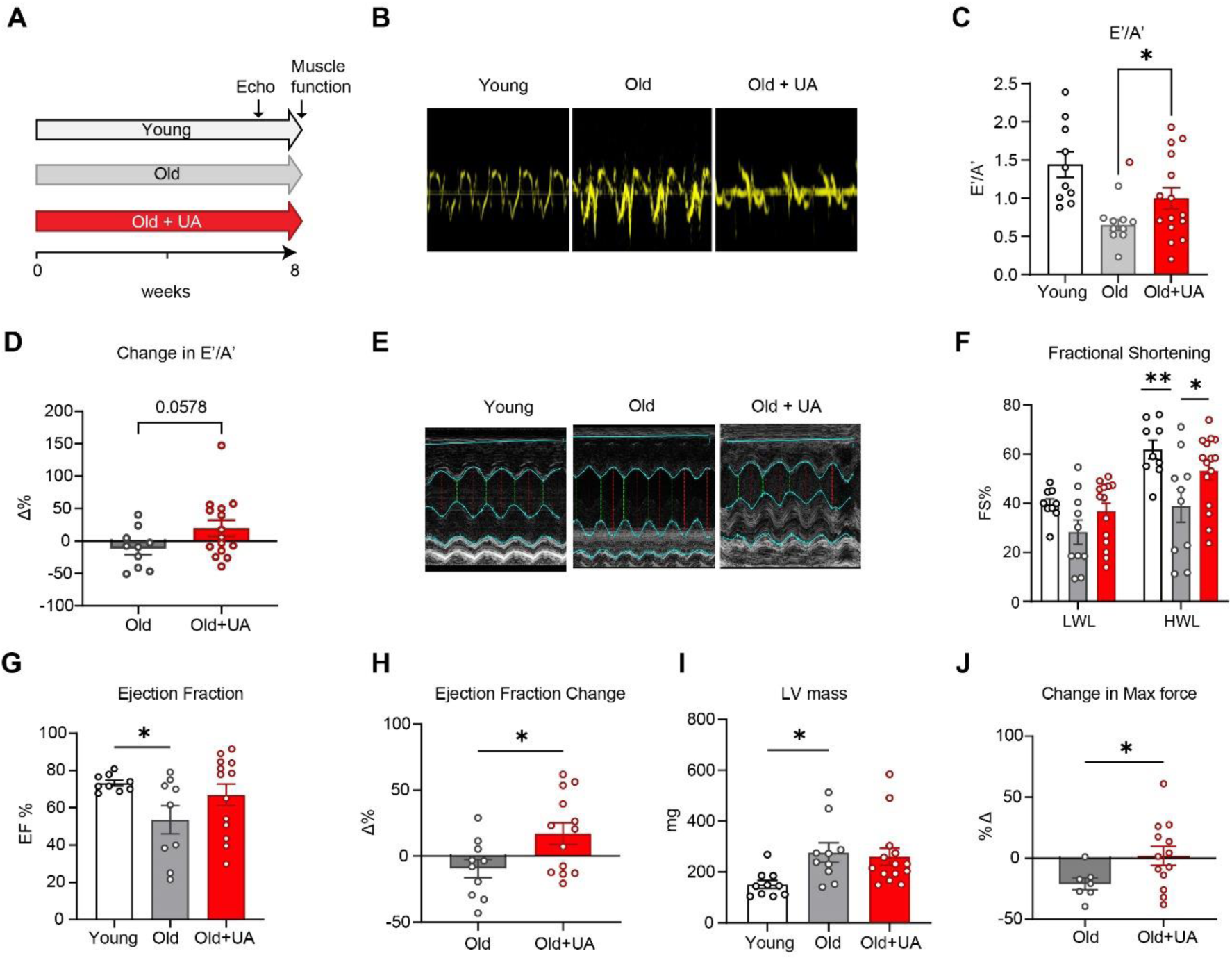
UA improves aging cardiac function and skeletal muscle force. A) Study schematics of both young and old mice feeding and experimental timeline. B) Representative image of echo Doppler flow measure used to assess diastolic function. C-D) Diastolic function of blood flow across the mitral valve in early diastole (Ea/E’) to the flow in late diastole (Aa/A’). Diastolic function represents early to late diastole ratio (E’/A’) in young and aged mice (C) and change in E’/A’ (post-treatment – baseline) in aged mice (D) E) Representative image of M mode short axis to assess left ventricle. F) Fractional shortening (FS%) under high and low workload. G-H) Ejection fraction at endpoint (G) and change in ejection fraction (H) in aged animals. I) Left ventricle mass from heart weights after sacrifice expressed in milligrams. J) Maximal torque of plantar flexor by nerve stimulation expressed as change over baseline. Animal replicates: Cardiac data: young=9, old=10, old + UA=15: Skeletal Muscle data: old=7, old+UA=13. * p<0.05; ** p<0.01 after t-test or post-hoc following one or two-way ANOVA. Error bars represent mean ± SEM.

### UA Improved Cardiac Function and Mitochondrial Health in a Model of HFrEF

Data above indicates a beneficial impact of UA on rescuing diastolic and systolic function in a model that partially recapitulates human HFpEF. We further determined the impact of Urolithin A on a rat model of heart failure with reduced ejection fraction (HFrEF). Myocardial infarction causing reduced systolic function was induced in Wistar Han rats by permanent ligation of the descending coronary artery (MI). A group undergoing sham surgery was used as control (Fig. 4A). As expected, MI rats showed a significant decrease in ejection fraction (EF) one week after surgery, compared to sham (Suppl. Fig. 4A). At this time point, MI animals were randomized based on their EF and administered by gavage with either UA at 25mpk (MI+UA) or vehicle (MI), for 2 months (Fig. 4A).

**Figure 4.**
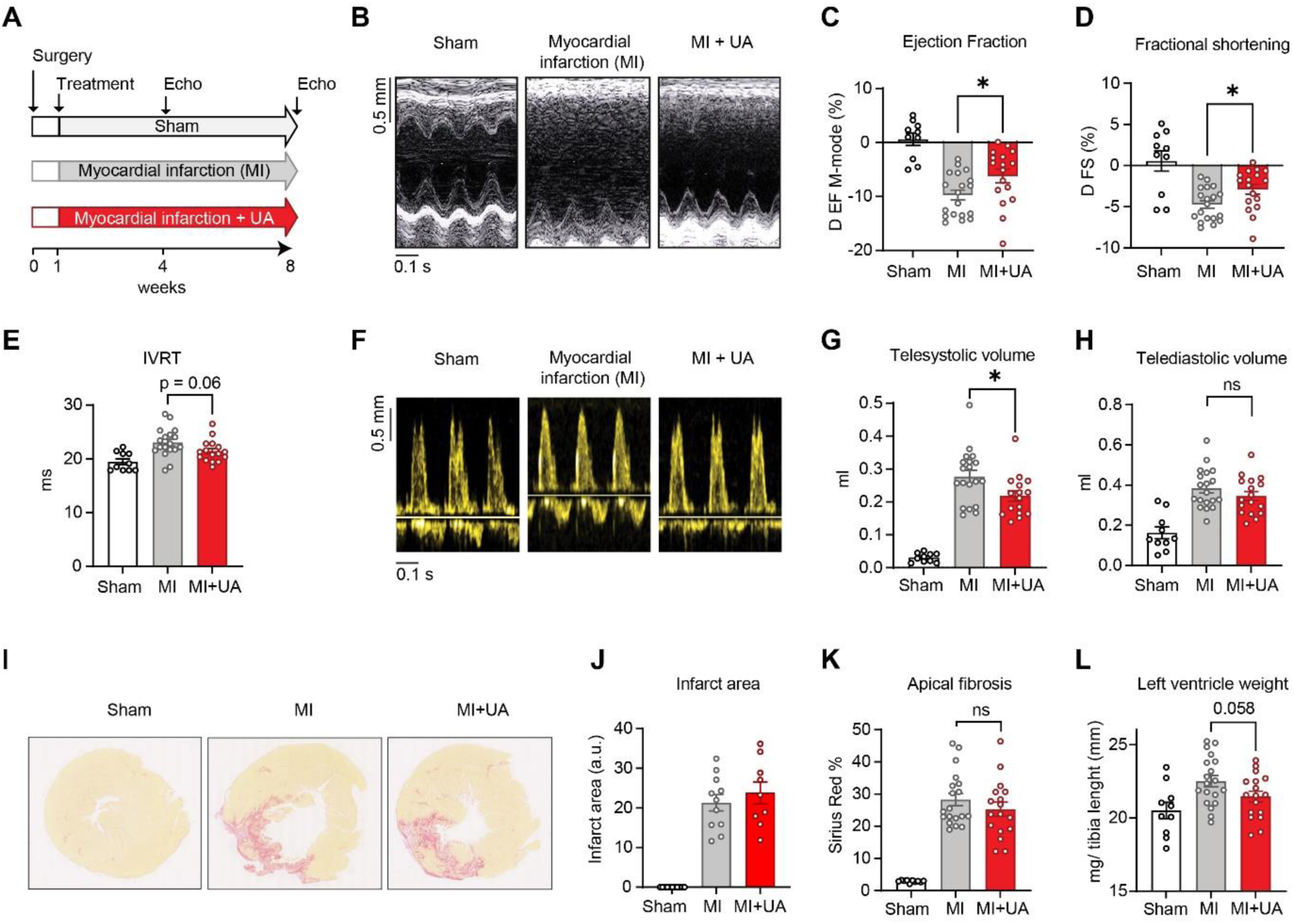
UA protects against cardiac dysfunction in a rat model of heart failure. A) Study schematics of surgery, treatments, and echocardiography timeline for control rats (Sham), rats with myocardial infarction (MI) and MI rats treated with Urolithin A (MI + UA). B) Representative images of M-mode measurement of the left ventricle structure used assess cardiac function and cardiac remodeling. C-D) Ejection fraction (EF) and fractional shortening (FS) measured as percentage change comparing end of the study to the start of the treatment (1 week time point) E) Measure of the isovolumic relaxation time (IVRT), as above F) Representative images of echo Doppler for Sham, MI and MI + UA group used to measure diastolic function. G-H) Telesystolic volume (G) and telediastolic volume (H) expressed as milliliters in the indicated groups at the end of the study. I-K) Representative images of left ventricle stained with Sirus-RED (I). Corresponding quantification of infarct area expressed as arbitrary units and fold change over sham (J) and of the apical fibrosis percentage expressed as percentage over total LV area (K). L) Measure of the left ventricle weight expressed as milligrams over tibia length at the end of the study in the indicated groups. Number of replicates: Sham (n=10), MI (n=19) and MI + UA (n=17). For figure E: Sham (n=10), MI (n=18) and MI + UA (17). For figure J, K: MI (n=11) and MI + UA (9) p value. *p<0.05; after one-way ANOVA. Error bars represent mean ± SEM.

Echocardiography revealed that UA administration for 2 months significantly improved systolic function in rats with heart failure. The decline of EF and FS, calculated as the delta between end to beginning of the treatment, was reduced by 35% and 39%, respectively, by comparing UA to vehicle groups (Fig. 4B-4D). Notably, a trend towards enhanced EF was observed with UA treatment already at the intermediate 1-month time point (Suppl. Fig. 4B). UA also showed a tendency to improve diastolic function, measured by Isovolumetric relaxation time (IVRT), despite this being a less striking feature in such a model of chronic heart failure (Fig. 4E and 4F).

Looking at cardiac remodeling parameters, telesystolic volume was dramatically increased by heart failure and significantly reduced following UA administration (Fig. 4G). This supports the beneficial impact of UA on cardiac muscle contractility. Other remodeling readouts, such as telediastolic volume (Fig. 4H) remained unchanged in UA versus sham groups. UA did not affect infarct size area (Fig. 4I and 4J) and apical ventricle fibrosis (Fig. 4K). Finally, a close to significant reduction in the ventricle weight was observed with UA treatment of MI mice (Fig. 4L). Altogether, these data indicate a cardioprotective action of UA administration in a model of HFrEF, with major impact of UA on systolic function restoration.

To investigate mechanisms underlying the in vivo cardioprotective effects of Urolithin A, we performed RNA-seq on heart sections from rats described above. A gene set enrichment analysis (GSEA) identified pathways that were significantly altered with HF and rescued by UA treatment, the majority of which were downregulated with HF and then increased by UA (Fig. 5A and Suppl. Table 3). They included several pathways associated with mitochondrial respiration, as well as nucleoid and peroxisome biology (Fig. 5B). A decline in mitochondrial gene sets in this rat model is consistent with data from human subjects (Fig.1B-D) and mouse aging study (Fig. 2E). Several genes whose expression is rescued by UA in diseased animals are related to mitochondrial OXPHOS complexes (Fig. 5C and Suppl. Fig. 4C). No changes were observed for mitophagy related genes in this setting. We therefore measured mitophagy activity in heart sections from the MI animals with or without UA treatment using the PINK1/Parkin-mediated mitophagy marker, phospho-Ubiquitin (ph-Ub). Consistent with previous studies in other age-related conditions^24,28,34^, UA was able increase ph-UB levels in the heart of MI animals, compared to vehicle (Fig. 5D and 5E).

**Figure 5.**
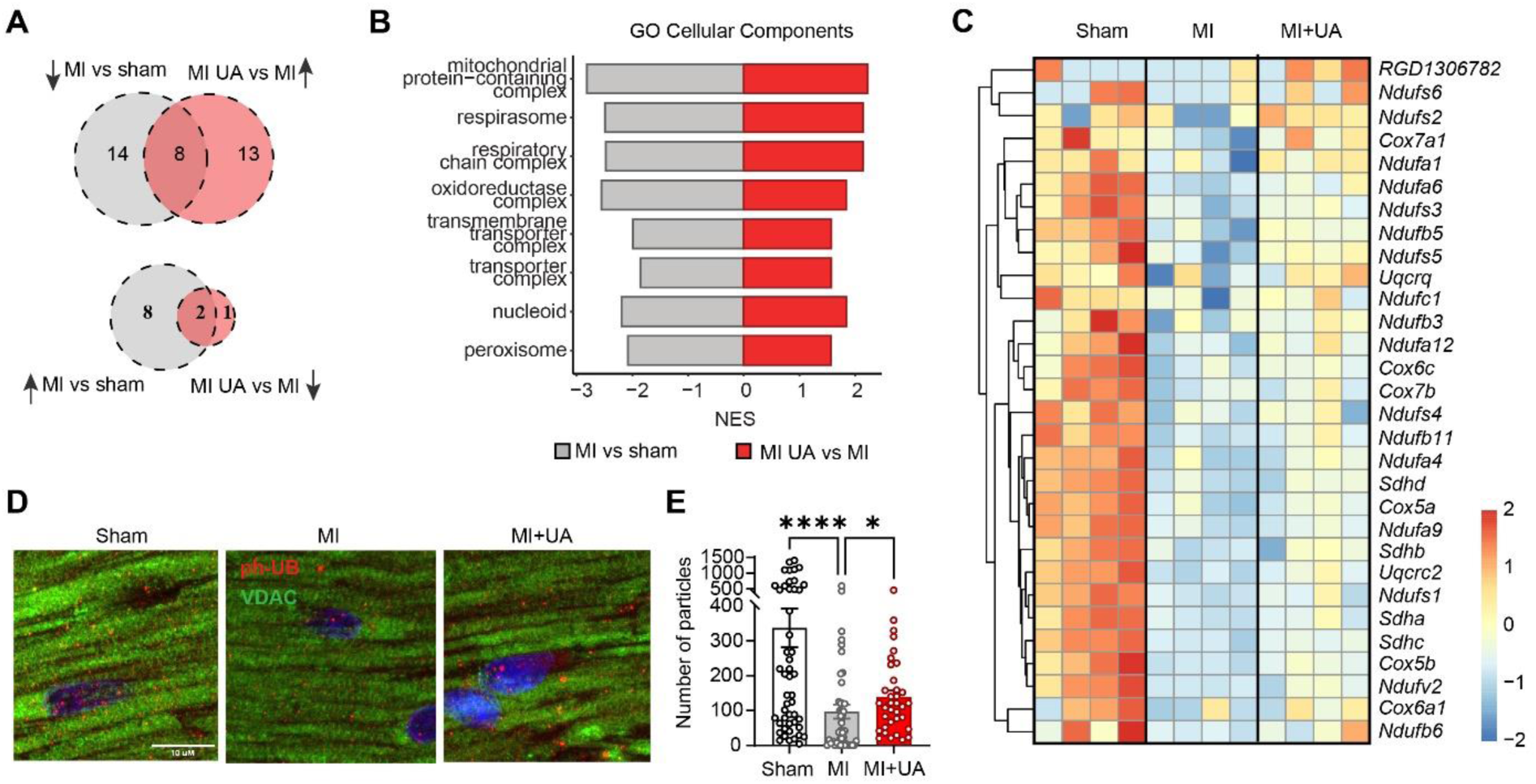
UA rescues defect in mitochondrial gene expression in rats with heart failure. A) Venn diagram depicting overlap of significantly enriched Gene Ontology Cellular Components (GO CCs) identified by Gene Set Enrichment Analysis (GSEA). Top diagram shows GO CCs suppressed in myocardial infarction (MI) versus sham and rescued by UA ("MI UA vs. MI") Lower panel: Similar as above, depicting GO CCs that are activated in the "MI vs. Sham" comparison and suppressed in the "MI UA vs. MI" comparison. B) Stacked bar graphs summarizing the 8 common GO CCs depicted in (A, Upper panel). The x-axis represents the Normalized Enrichment Score (NES), indicating the level of suppression (negative values) or activation (positive values). The ‘MI vs. Sham’ comparison is represented in gray, while the ‘MI UA vs. MI’ comparison is shown in red. C) Heatmaps displaying scaled normalized expression counts (z-scores) of the genes that contributed to the enrichment of the "Respiratory Chain Complex " GO CC term in both “MI vs Sham” and “MI UA vs MI” comparisons (respectively suppressed and activated). Rows represent genes and were hierarchically clustered using a Euclidean distance metric and complete linkage. Columns represent individual samples from each condition. (n = 4). F-G) Representative images of left ventricles (LV) stained for VDAC (green) and phospho-ubiquitin (ph-UB, red) in the indicated groups. Nuclei were stained in blue with DAPI (f). Quantification of the number of particles corresponding to ph-Ub staining (G). (N = 54 images/group) *p value: ****p < 0.001 after one-way ANOVA. Error bars represent mean ± SEM.

## Discussion

Heart diseases are world’s main cause of death causing 17.9 million deaths worldwide^35^. The geroscience approach recognizes that aging is the major risk factor for most chronic diseases, including heart disease, and by developing approaches to target fundamental aspects of aging, we can reduce the disease burden of the most common chronic diseases^36^. Mitochondrial dysfunction, one of the twelve hallmarks of aging^37^, has also been shown to be a mechanistic driver of heart failure in multiple pre-clinical models^38,39,40^.

Our unique comparison of gene expression changes in the aging human heart from the GTEx study^32^ with the transcriptional profiles from multiple cardiomyopathies in humans^33^ supports a focus on mitochondrial targeted interventions for both aging and heart disease. We identified mitochondrial genes, particularly those associated with oxidative phosphorylation, mitochondrial quality control, and mitophagy, among the most prominent transcriptional responses to aging, dilated and ischemic cardiomyopathy. This intersection of cardiac aging and cardiomyopathies in the human heart points to the translational relevance of improving mitochondrial quality control to protect against the onset and progression of heart dysfunction.

Indeed, we showed that a mitophagy enhancer, urolithin A, significantly counteracts mitochondrial and heart dysfunctions in preclinical models of both cardiac aging, typified by preserved ejection fraction, and heart failure characterized by reduced ejection fraction. Previous work has demonstrated that enhancing mitophagy by treating with UA can reduces cardiac dysfunction in models of diabetes^34,41^, obesity^42^ and sepsis^43^. The present report is the first investigation of the beneficial effects of UA on the aging heart and builds on both preclinical^24^ and clinical studies^27,28^ demonstrating positive impact of UA treatment on aging skeletal muscle. The decline in mitochondrial quality control in the aging mouse hearts was supported by the disruption in mitochondrial cristae structure, consistent with previous reports from both age and heart failure models^44,45,46^, despite no overall decline in mitochondrial volume density. Rescuing mitochondrial quality control with 8 weeks of oral gavage UA treatment reversed the disrupted ultrastructure and mitochondrial gene expression changes associated with aging, while no effect on mitochondrial respiration was observed in a parallel study after 8 weeks of UA by food admix. It is noteworthy that the mitolysosome structures in the aged UA treated mice significantly overshot those in the young and young UA-treated hearts, suggesting that UA does not just enhance mitophagy, but boosts the sensitivity of the mitophagic system to respond to poorly functioning mitochondria.

Muscle fueling through mitochondria fatty acid oxidation is essential to provide enough cellular energy. In the skeletal muscle this is reflected by a tight crosstalk and energy transfer between lipid droplets and mitochondria^47^. The impact of this crosstalk on the cardiac muscle was not elucidated. Here, we show a marked lipid droplet volume density depletion during cardiac aging and a significant replenishment after UA supplementation. This supports further research on heart lipid-mitochondria interactions and identified a novel potential mechanism by which UA improves cellular bioenergetics.

A key feature of the aging mouse and human heart is diastolic dysfunction, or heart failure with preserved ejection fraction (HFpEF)^48,49,50^. As a result, many previous studies with mitochondrial interventions focused primarily on improvements in diastolic dysfunction^18,51,52^. In this study we observed systolic ejection fraction and fractional shortening, as well as diastolic deficits in the aging mouse heart that were partially reversed by UA treatment for 8 weeks in the chow. The effects of age and UA were enhanced by stimulating cardiac work with dobutamine to a more physiologically relevant workload. These results make UA unique among a growing list of mitochondrial interventions that can restore function in the aging heart, in that UA appears to improve both diastolic and systolic function. Other interventions focused on enhancing autophagy/mitophagy like spermidine were reported to improve diastolic dysfunction^51^, while interventions targeting NAD metabolism like NMN improved systolic dysfunction^53^, alone. Elamipretide was initially reported to only improve diastolic dysfunction in the aging heart^54^, but more recent work using the more sensitive echocardiography with global longitudinal strain analysis demonstrates significant effects on systolic function as well^55^.

Despite the focus on HFpEF with age, many age-related changes in the heart increase the susceptibility of the aging heart to heart failure with reduced ejection fraction (HFrEF). We show the ability of UA supplementation to improve mitochondrial and cardiac function in a rat myocardial ischemia model of chronic heart failure. In this model, urolithin A significantly improved systolic function and rescued gene expression changes observed in diseased rats after 8-weeks of administration^56^.

Exercise and adequate intake of Omega-3 are among the top recommended strategies. Indeed, exercise is a Class 1 recommendation for patients with HF^57^ and the American Heart Association promotes performing regular exercise to support cardiovascular health^56^. Interestrigly, an endurance exercise regimen of the same 8-weeks duration in a HF rat model, exerted similar benefits as UA on heart function and mitochondrial bioenergetics^58^. Adequate omega-3 fatty acids consumption has been associated with significant reduction in the risk of coronary heart disease. Mechanisms underlying the benefits of omega-3 for heart health includes lowering triglycerides and increasing healthy high-density lipoprotein levels^59^. Notably, this is a potentially complimentary approach to increasing mitochondrial quality by UA. Therefore, our data support the potential of UA to support HFrEF patients and to maintain optimal cardiovascular function throughout life, alone and in combination with omega-3 intake and exercise regimens.

### Clinical Relevance

Urolithin A is a naturally occurring compound that is converted by the gut microbiome in humans^23^. Only 30% to 40% of the adult population has the correct microbiome composition to produce UA and even in those with the correct microbiome composition achieving levels of UA used in this study is not practical through normal dietary intake^60^. Therefore, direct UA supplementation may be an important tool to improve quality of life with age. Urolithin A has already been tested in clinical studies with proven safety and positive clinical endpoints related to improved muscle function. In older adults two months of supplementation led to increased muscle endurance^27^. In middle aged overweight subjects, 4 months of UA supplementation increased VO2 by 10%^28^. Our data suggests that improved heart function might contribute to the positive increase of muscle function and especially VO2max measured physical performance.

In addition, we highlight that urolithin A supplementation showed maintenance of skeletal muscle force in aged mice. This is clinically relevant as skeletal muscle mitochondrial function is impaired in patients with HFpEF and contributes to the reduced exercise tolerance and quality of life. The ability of UA to improve both cardiac and skeletal muscle function could be used to facilitate the development and validation of therapeutic strategies of mitochondrial booster to combat HFpEF^61^,and supports UA’s potential for managing heart and skeletal muscle health throughout life.

### Limitations

The main limitation of this study is that the data from the aging heart comes from two parallel studies, one where the UA was delivered by daily oral gavage and used to assess mitochondrial ultrastructure and gene expression and the other where UA was provided in the chow used for in vivo cardiac functional assays. Ideally both mitochondria ultrastructure and in vivo function would have been measured in both studies. However, given the different dosing strategies and endpoints in the two studies it is noteworthy that both delivery methods yielded significant reversal of aspects of cardiac aging phenotypes providing further evidence for a robust and reproducible effect of UA on the aging heart. A further limitation of the heart failure data is that they do not capture the effect of UA on acute HF. Additional studies are needed to address whether improving cardiac health with UA treatment can increase resilience and preserve function following acute cardiac stress such as ischemia reperfusion.

### Conclusion

We demonstrate significant improvement in cardiac function associated with improved mitochondrial quality control following treatment with UA in two heart dysfunction models, the aging mouse heart and myocardial infarction in rats. Together with previous results from clinical trials demonstrating improved muscle function, these data point to UA supplementation as a tool to enhance healthy aging and quality of life in older adults and call for clinical studies validating the impact of UA on cardiac function in humans.

## Data availability

RNA-seq data that support the findings of this study will be deposited in the NCBI Gene Expression Omnibus (GEO; http://www.ncbi.nlm.nih.gov/geo/).

## Acknowledgement

We thank Leonidas Karagounis Global Science Lead, Muscle Metabolism & Metabolic Health at Nestlé Heath Science during the course of this study and now Professor of Research Translation & Enterprise Professor of Research Translation & Enterprise at the Australian Catholic University for the support in his critical comments during the design of the studies.

## Authors Contributions

D.D., D.J.M, J.F. and P.A.A contributed to the design of the study. J.F. coordinated the cardiac aging study and collected samples for Electron Microscopy (EM) (Study 1). R.L and T.D performed Electron Microscopy imaging. J.F. and C.T. analysed EM images and interpreted data. S.L. performed the cardiac aging study 2 and interpreted all data; G.B. performed the rat heart failure study. C.T. performed all immunofluorescence analyses and J.F. and C.T. interpreted these data. C.T. performed sample preparation for RNA-seq analyses. D.K. performed all bioinformatic analyses. D.D. and D.J.M. wrote the manuscript, with the help of the other co-authors. All authors reviewed the manuscript.

## Supplemental Material

Supplemental Figures S1-S4

Supplemental Tables S1-S3

## References

1. Birger M, Kaldjian AS, Roth GA, Moran AE, Dieleman JL, Bellows BK. Spending on Cardiovascular Disease and Cardiovascular Risk Factors in the United States: 1996 to 2016. Circulation. 2021;144(4):271–282.

2. Yazdanyar A, Newman AB. The burden of cardiovascular disease in the elderly: morbidity, mortality, and costs. Clinics in Geriatric Medicine. 2009;25(4):563–577, vii.

3. Paneni F, Diaz Cañestro C, Libby P, Lüscher TF, Camici GG. The Aging Cardiovascular System: Understanding It at the Cellular and Clinical Levels. Journal of the American College of Cardiology. 2017;69(15):1952–1967.

4. Barasch E, Gottdiener JS, Aurigemma G, Kitzman DW, Han J, Kop WJ, Tracy RP. Association between elevated fibrosis markers and heart failure in the elderly: the cardiovascular health study. Circulation. Heart Failure. 2009;2(4):303–310.

5. Rondelli RR, Dal Corso S, Simões A, Malaguti C. Methods for the assessment of peripheral muscle fatigue and its energy and metabolic determinants in COPD. Jornal Brasileiro De Pneumologia: Publicacao Oficial Da Sociedade Brasileira De Pneumologia E Tisilogia. 2009;35(11):1125–1135.

6. Smart N, Haluska B, Jeffriess L, Marwick TH. Exercise training in systolic and diastolic dysfunction: effects on cardiac function, functional capacity, and quality of life. American Heart Journal. 2007;153(4):530–536.

7. Williams S, Wynn G, Cozza K, Sandson NB. Cardiovascular medications. Psychosomatics. 2007;48(6):537–547.

8. Roger VL, Go AS, Lloyd-Jones DM, Benjamin EJ, Berry JD, Borden WB, Bravata DM, Dai S, Ford ES, Fox CS, Fullerton HJ, Gillespie C, Hailpern SM, Heit JA, Howard VJ, et al. Heart disease and stroke statistics--2012 update: a report from the American Heart Association. Circulation. 2012;125(1):e2–e220.

9. Parikh JD, Hollingsworth KG, Wallace D, Blamire AM, MacGowan GA. Normal age-related changes in left ventricular function: Role of afterload and subendocardial dysfunction. International Journal of Cardiology. 2016;223:306–312.

10. Parikh JD, Hollingsworth KG, Wallace D, Blamire AM, MacGowan GA. Left ventricular functional, structural and energetic effects of normal aging: Comparison with hypertension. PLoS ONE. 2017;12(5):e0177404.

11. Haykowsky MJ, Timmons MP, Kruger C, McNeely M, Taylor DA, Clark AM. Meta-analysis of aerobic interval training on exercise capacity and systolic function in patients with heart failure and reduced ejection fractions. The American Journal of Cardiology. 2013;111(10):1466–1469.

12. Haykowsky MJ, Kitzman DW. Exercise physiology in heart failure and preserved ejection fraction. Heart Failure Clinics. 2014;10(3):445–452.

13. Dai D-F, Chiao YA, Marcinek DJ, Szeto HH, Rabinovitch PS. Mitochondrial oxidative stress in aging and healthspan. Longevity & Healthspan. 2014;3:6.

14. Dai D-F, Santana LF, Vermulst M, Tomazela DM, Emond MJ, MacCoss MJ, Gollahon K, Martin GM, Loeb LA, Ladiges WC, Rabinovitch PS. Overexpression of catalase targeted to mitochondria attenuates murine cardiac aging. Circulation. 2009;119(21):2789.

15. Brown DA, Perry JB, Allen ME, Sabbah HN, Stauffer BL, Shaikh SR, Cleland JGF, Colucci WS, Butler J, Voors AA, Anker SD, Pitt B, Pieske B, Filippatos G, Greene SJ, et al. Mitochondrial function as a therapeutic target in heart failure. Nature reviews. Cardiology. 2017;14(4):238–250.

16. Opie LH. Review of trials in the treatment of coronary artery disease: theoretical expectations versus lack of practical success--how can we explain the differences? The American Journal of Cardiology. 1998;82(3A):15H–20H.

17. Andersson DC, Marks AR. Fixing ryanodine receptor Ca2+ leak - a novel therapeutic strategy for contractile failure in heart and skeletal muscle. Drug discovery today. Disease mechanisms. 2010;7(2):e151–e157.

18. Chiao YA, Zhang H, Sweetwyne M, Whitson J, Ting YS, Basisty N, Pino LK, Quarles E, Nguyen N-H, Campbell MD, Zhang T, Gaffrey MJ, Merrihew G, Wang L, Yue Y, et al. Late-life restoration of mitochondrial function reverses cardiac dysfunction in old mice. eLife. 9:e55513.

19. Dai D-F, Rabinovitch P. Mitochondrial oxidative stress mediates induction of autophagy and hypertrophy in angiotensin-II treated mouse hearts. Autophagy. 2011;7(8):917–918.

20. Hegland A. Just passing through. This innovative evaluation unit stresses teamwork to place patients in less-costly facilities or return them to their homes. Contemporary Longterm Care. 1994;17(4):61–62.

21. Wen J, Pan T, Li H, Fan H, Liu J, Cai Z, Zhao B. Role of mitophagy in the hallmarks of aging. Journal of Biomedical Research. 2022;37(1):1–14.

22. Guo J, Chiang W-C. Mitophagy in aging and longevity. IUBMB life. 2022;74(4):296–316.

23. D’Amico D, Andreux PA, Valdés P, Singh A, Rinsch C, Auwerx J. Impact of the Natural Compound Urolithin A on Health, Disease, and Aging. Trends in Molecular Medicine. 2021;27(7):687–699.

24. Ryu D, Mouchiroud L, Andreux PA, Katsyuba E, Moullan N, Nicolet-Dit-Félix AA, Williams EG, Jha P, Lo Sasso G, Huzard D, Aebischer P, Sandi C, Rinsch C, Auwerx J. Urolithin A induces mitophagy and prolongs lifespan in C. elegans and increases muscle function in rodents. Nature Medicine. 2016;22(8):879–888.

25. Luan P, D’Amico D, Andreux PA, Laurila P-P, Wohlwend M, Li H, Lima TI de, Place N, Rinsch C, Zanou N, Auwerx J. Urolithin A improves muscle function by inducing mitophagy in muscular dystrophy. Science Translational Medicine. 2021;13(588).

26. D’Amico D, Olmer M, Fouassier AM, Valdés P, Andreux PA, Rinsch C, Lotz M. Urolithin A improves mitochondrial health, reduces cartilage degeneration, and alleviates pain in osteoarthritis. Aging Cell. 2022;21(8):e13662.

27. Liu S, D’Amico D, Shankland E, Bhayana S, Garcia JM, Aebischer P, Rinsch C, Singh A, Marcinek DJ. Effect of Urolithin A Supplementation on Muscle Endurance and Mitochondrial Health in Older Adults: A Randomized Clinical Trial. JAMA network open. 2022;5(1):e2144279.

28. Singh A, D’Amico D, Andreux PA, Fouassier AM, Blanco-Bose W, Evans M, Aebischer P, Auwerx J, Rinsch C. Urolithin A improves muscle strength, exercise performance, and biomarkers of mitochondrial health in a randomized trial in middle-aged adults. Cell Reports Medicine. 2022;3(5).

29. Yang J, Huang T, Petralia F, Long Q, Zhang B, Argmann C, Zhao Y, Mobbs CV, Schadt EE, Zhu J, Tu Z, GTEx Consortium. Synchronized age-related gene expression changes across multiple tissues in human and the link to complex diseases. Scientific Reports. 2015;5:15145.

30. Siegel MP, Kruse SE, Percival JM, Goh J, White CC, Hopkins HC, Kavanagh TJ, Szeto HH, Rabinovitch PS, Marcinek DJ. Mitochondrial-targeted peptide rapidly improves mitochondrial energetics and skeletal muscle performance in aged mice. Aging Cell. 2013;12(5):763–771.

31. Wolf DM, Segawa M, Kondadi AK, Anand R, Bailey ST, Reichert AS, van der Bliek AM, Shackelford DB, Liesa M, Shirihai OS. Individual cristae within the same mitochondrion display different membrane potentials and are functionally independent. The EMBO journal. 2019;38(22):e101056.

32. GTEx Consortium. The Genotype-Tissue Expression (GTEx) project. Nature Genetics. 2013;45(6):580–585.

33. Sweet ME, Cocciolo A, Slavov D, Jones KL, Sweet JR, Graw SL, Reece TB, Ambardekar AV, Bristow MR, Mestroni L, Taylor MRG. Transcriptome analysis of human heart failure reveals dysregulated cell adhesion in dilated cardiomyopathy and activated immune pathways in ischemic heart failure. BMC genomics. 2018;19(1):812.

34. Huang C, Huang L, Huang Q, Lin L, Wang L, Wu Y, Wu K, Gao R, Liu X, Liu X, Qi L, Liu L. Mitophagy disorder mediates cardiac deterioration induced by severe hypoglycemia in diabetic mice. Molecular and Cellular Endocrinology. 2023:111994.

35. Anon. Cardiovascular diseases.

36. Bartke A. New Directions in Research on Aging. Stem Cell Reviews and Reports. 2022;18(4):1227–1233.

37. López-Otín C, Blasco MA, Partridge L, Serrano M, Kroemer G. Hallmarks of aging: An expanding universe. Cell. 2023;186(2):243–278.

38. Sabbah HN, Gupta RC, Kohli S, Wang M, Hachem S, Zhang K. Chronic Therapy With Elamipretide (MTP-131), a Novel Mitochondria-Targeting Peptide, Improves Left Ventricular and Mitochondrial Function in Dogs With Advanced Heart Failure. Circulation. Heart Failure. 2016;9(2):e002206.

39. Karamanlidis G, Lee CF, Garcia-Menendez L, Kolwicz SC, Suthammarak W, Gong G, Sedensky MM, Morgan PG, Wang W, Tian R. Mitochondrial complex I deficiency increases protein acetylation and accelerates heart failure. Cell Metabolism. 2013;18(2):239–250.

40. Caudal A, Tang X, Chavez JD, Keller A, Mohr JP, Bakhtina AA, Villet O, Chen H, Zhou B, Walker MA, Tian R, Bruce JE. Mitochondrial interactome quantitation reveals structural changes in metabolic machinery in the failing murine heart. Nature Cardiovascular Research. 2022;1(9):855–866.

41. Savi M, Bocchi L, Mena P, Dall’Asta M, Crozier A, Brighenti F, Stilli D, Del Rio D. In vivo administration of urolithin A and B prevents the occurrence of cardiac dysfunction in streptozotocin-induced diabetic rats. Cardiovascular Diabetology. 2017;16(1):80.

42. Huang J-R, Zhang M-H, Chen Y-J, Sun Y-L, Gao Z-M, Li Z-J, Zhang G-P, Qin Y, Dai X- Y, Yu X-Y, Wu X-Q. Urolithin A ameliorates obesity-induced metabolic cardiomyopathy in mice via mitophagy activation. Acta Pharmacologica Sinica. 2023;44(2):321–331.

43. Wang Y, Jasper H, Toan S, Muid D, Chang X, Zhou H. Mitophagy coordinates the mitochondrial unfolded protein response to attenuate inflammation-mediated myocardial injury. Redox Biology. 2021;45:102049.

44. Allen ME, Pennington ER, Perry JB, Dadoo S, Makrecka-Kuka M, Dambrova M, Moukdar F, Patel HD, Han X, Kidd GK, Benson EK, Raisch TB, Poelzing S, Brown DA, Shaikh SR. The cardiolipin-binding peptide elamipretide mitigates fragmentation of cristae networks following cardiac ischemia reperfusion in rats. Communications Biology. 2020;3(1):389.

45. Morris S, Molina-Riquelme I, Barrientos G, Bravo F, Aedo G, Gómez W, Lagos D, Verdejo H, Peischard S, Seebohm G, Psathaki OE, Eisner V, Busch KB. Inner mitochondrial membrane structure and fusion dynamics are altered in senescent human iPSC-derived and primary rat cardiomyocytes. Biochimica Et Biophysica Acta. Bioenergetics. 2023;1864(2):148949.

46. Gong Y, Luo Y, Liu S, Ma J, Liu F, Fang Y, Cao F, Wang L, Pei Z, Ren J. Pentacyclic triterpene oleanolic acid protects against cardiac aging through regulation of mitophagy and mitochondrial integrity. Biochimica Et Biophysica Acta. Molecular Basis of Disease. 2022;1868(7):166402.

47. Parry HA, Glancy B. Energy transfer between the mitochondrial network and lipid droplets in insulin resistant skeletal muscle. Current opinion in physiology. 2021;24:100487.

48. Obokata M, Sorimachi H, Harada T, Kagami K, Saito Y, Ishii H. Epidemiology, Pathophysiology, Diagnosis, and Therapy of Heart Failure With Preserved Ejection Fraction in Japan. Journal of Cardiac Failure. 2023;29(3):375–388.

49. Hellon CP, Solomon MI. Suicide and age in Alberta, Canada, 1951 to 1977. The changing profile. Archives of General Psychiatry. 1980;37(5):505–510.

50. Chiao YA, Rabinovitch PS. The Aging Heart. Cold Spring Harbor Perspectives in Medicine. 2015;5(9):a025148.

51. Eisenberg T, Abdellatif M, Schroeder S, Primessnig U, Stekovic S, Pendl T, Harger A, Schipke J, Zimmermann A, Schmidt A, Tong M, Ruckenstuhl C, Dammbrueck C, Gross AS, Herbst V, et al. Cardioprotection and lifespan extension by the natural polyamine spermidine. Nature Medicine. 2016;22(12):1428–1438.

52. Quarles E, Basisty N, Chiao YA, Merrihew G, Gu H, Sweetwyne MT, Fredrickson J, Nguyen N-H, Razumova M, Kooiker K, Moussavi-Harami F, Regnier M, Quarles C, MacCoss M, Rabinovitch PS. Rapamycin persistently improves cardiac function in aged, male and female mice, even following cessation of treatment. Aging Cell. 2020;19(2):e13086.

53. Whitson JA, Bitto A, Zhang H, Sweetwyne MT, Coig R, Bhayana S, Shankland EG, Wang L, Bammler TK, Mills KF, Imai S-I, Conley KE, Marcinek DJ, Rabinovitch PS. SS-31 and NMN: Two paths to improve metabolism and function in aged hearts. Aging Cell. 2020;19(10):e13213.

54. Chiao YA, Zhang H, Sweetwyne M, Whitson J, Ting YS, Basisty N, Pino LK, Quarles E, Nguyen N-H, Campbell MD, Zhang T, Gaffrey MJ, Merrihew G, Wang L, Yue Y, et al. Late-life restoration of mitochondrial function reverses cardiac dysfunction in old mice. eLife. 2020;9:e55513.

55. Pharaoh G, Kamat V, Kannan S, Stuppard RS, Whitson J, Martin-Perez M, Qian W-J, MacCoss MJ, Villen J, Rabinovitch P, Campbell MD, Sweet IR, Marcinek DJ. Elamipretide Improves ADP Sensitivity in Aged Mitochondria by Increasing Uptake through the Adenine Nucleotide Translocator (ANT). 2023:2023.02.01.525989.

56. Anon. How much physical activity do you need? www.heart.org.

57. Jl T, J M, Ar B. Practical guidelines for exercise prescription in patients with chronic heart failure. Heart failure reviews. 2023.

58. J K, J M, D P, P Z, Z D, U W, M L. Aerobic interval training attenuates remodelling and mitochondrial dysfunction in the post-infarction failing rat heart. Cardiovascular research. 2013;99(1).

59. Chaddha A, Eagle KA. Omega-3 Fatty Acids and Heart Health. Circulation. 2015;132(22):e350–e352.

60. Singh A, D’Amico D, Andreux PA, Dunngalvin G, Kern T, Blanco-Bose W, Auwerx J, Aebischer P, Rinsch C. Direct supplementation with Urolithin A overcomes limitations of dietary exposure and gut microbiome variability in healthy adults to achieve consistent levels across the population. European Journal of Clinical Nutrition. 2022;76(2):297–308.

61. Scandalis L, Kitzman DW, Nicklas BJ, Lyles M, Brubaker P, Nelson MB, Gordon M, Stone J, Bergstrom J, Neufer PD, Gnaiger E, Molina AJA. Skeletal Muscle Mitochondrial Respiration and Exercise Intolerance in Patients With Heart Failure With Preserved Ejection Fraction. JAMA Cardiology. 2023.

